# Protective Effects of STI-2020 Antibody Delivered Post-Infection by the Intranasal or Intravenous Route in a Syrian Golden Hamster COVID-19 Model

**DOI:** 10.1101/2020.10.28.359836

**Authors:** Yanwen Fu, Junki Maruyama, Alok Singh, Reyna Lim, Arthur Ledesma, Daniel Lee, Laura Rivero-Nava, Jamie Ko, Ianne Rivera, Rachel A. Sattler, John T. Manning, Lisa Kerwin, Heyue Zhou, Mark Brunswick, Damien Bresson, Henry Ji, Slobodan Paessler, Robert D. Allen

**Author notes:** these authors contributed equally to the work (Yanwen Fu, Junki Maruyama, Alok Singh).

## Abstract

We have previously reported that the SARS-CoV-2 neutralizing antibody, STI-2020, potently inhibits cytopathic effects of infection by genetically diverse clinical SARS-CoV-2 pandemic isolates in vitro, and has demonstrated efficacy in a hamster model of COVID-19 when administered by the intravenous route immediately following infection. We now have extended our in vivo studies of STI-2020 to include disease treatment efficacy, profiling of biodistribution of STI-2020 in mice when antibody is delivered intranasally (IN) or intravenously (IV), as well as pharmacokinetics in mice following IN antibody administration. Importantly, SARS-CoV-2-infected hamsters were treated with STI-2020 using these routes, and treatment effects on severity and duration of COVID-19-like disease in this model were evaluated. In SARS-CoV-2 infected hamsters, treatment with STI-2020 12 hours post-infection using the IN route led to a decrease in severity of clinical disease signs and a more robust recovery during 9 days of infection as compared to animals treated with an isotype control antibody. Treatment via the IV route using the same dose and timing regimen resulted in a decrease in the average number of consecutive days that infected animals experienced weight loss, shortening the duration of disease and allowing recovery to begin more rapidly in STI-2020 treated animals. Following IN administration in mice, STI-2020 was detected within 10 minutes in both lung tissue and lung lavage. The half-life of STI-2020 in lung tissue is approximately 25 hours. We are currently investigating the minimum effective dose of IN-delivered STI-2020 in the hamster model as well as establishing the relative benefit of delivering neutralizing antibodies by both IV and IN routes.

## INTRODUCTION

The SARS-CoV-2 neutralizing monoclonal antibody (mAb) STI-2020 is an affinity-matured variant of the parental antibody STI-1499, first isolated from the fully human G-MAB phage display antibody library and paired with a human IgG1 Fc domain bearing the double mutation in the hinge region, L234A, L235A (LALA). This modification is aimed at reducing the potential for antibody-dependent enhancement (ADE) of infection in the context of COVID-19 treatment^1, 2, 3, 4, 5^. STI-2020 administered IV immediately following infection has been previously shown to provide protection against pathogenesis in the hamster COVID-19 disease model^5^. The hamster model provides a ready means of evaluating the effects of candidate therapeutics on SARS-CoV-2-mediated respiratory disease, which has contributed greatly to the overall morbidity and mortality associated with the current pandemic^6, 7, 8^. Human SARS-CoV-2 infection along the respiratory tract has been detected in the ciliated epithelial cells of the trachea, alveolar cells, and upper airway epithelia in tissues from COVID-19 autopsies^9, 10^. Expression of angiotensin converting enzyme-2 (ACE2) and neuropilin-1 (NRP1), both of which have been identified as entry factors mediating uptake of SARS-CoV-2 into host cells, has been detected at varying levels in the epithelia of the upper and lower respiratory tract ^9, 10, 11, 12^. Establishment and maintenance of a neutralizing antibody (nAb) blockade in regions of the respiratory tract that are most closely linked to primary virus infection, virus receptor expression, and progressive virus pathogenesis could provide a means of decreasing the severity of COVID-19 symptoms, preventing the dissemination of disease within the respiratory tract, and decreasing pharyngeal shedding of virus into the environment. Previously, it was demonstrated that mAbs directed against the influenza hemagglutinin (HA) stalk provide protection and therapeutic benefit to infected mice when administered as an aerosol or as a nasal droplet^13^. Recent studies have demonstrated the beneficial effects of nebulizer-based delivery of SARS-CoV-2 nAbs on severity of lung pathology and degree of virus replication in the lungs of infected hamsters^14^. To investigate alternative routes and timings of antibody administration, an intranasal STI-2020 formulation was developed for use in biodistribution, pharmacokinetic, and virus neutralization efficacy profiling experiments. Intranasally-delivered STI-2020 was detectable in mouse lung lavage and lung tissue within 10 minutes of administration. Following IN administration of a single 500 μg dose of STI-2020 twelve hours post-infection in the Syrian golden hamster COVID-19 model, animals exhibited decreased weight loss as compared to isotype control-treated animals and underwent a more rapid and robust recovery from disease than animals in the control treatment group. Therapeutic effects including decreased duration of progressive disease following treatment with STI-2020 were also observed in animals administered an equivalent dose IV at 12 hours post-infection.

## MATERIALS AND METHODS

### Formulation of STI-2020 monoclonal antibody

STI-2020 was produced, purified, and formulated as previously described^5^. For IN dosing of STI-2020, antibody was formulated in IN formulation buffer (20mM Histidine-HCl (cat# 4395-06, Macron), 240 mM Sucrose (cat# 4074-01, J.T. Baker), 0.05% Polysorbate 80 (cat# HX2, NOF Corporation), 0.3% Hydroxypropyl Methyl Cellulose (HPMC) (cat# H1335, Spectrum Chemical), pH 5.8).

### Antibody Detection ELISA

Multi-Array 96-well plates (cat# L15XA-3, Meso Scale Discovery (MSD)) were coated with mouse anti-human IgG antibody (CH2 domain, cat# MA5-16929, ThermoFisher Scientific) at 2 μg/mL in 1X PBS (50μL/well), sealed, and incubated overnight at 4°C. The following day, plates were washed 3X with 1X washing solution (KPL wash solution, cat no# 5150-0009, lot no# 10388555, Sera Care). Plates were then blocked using 50 μL/well of Blocker™ Casein in PBS (cat no# 37528, lot# QE220946, ThermoFisher) for 1 hour at room temperature on an orbital shaker. Plates were washed 3X with 1X washing solution. Samples from biodistribution or pharmacokinetic experiments were added in a volume of 50 μL to each well. STI-2020 antibody was serially diluted from a concentration of 1000 ng/mL to 3.1 ng/mL in Blocker™ Casein in PBS to generate the standard curve for the assay. Following addition of experimental samples or control samples, plates were incubated for 2 hours at room temperature on an orbital shaker. Plates were then washed 3X with 1X washing solution and 50 μL of Sulfo-Tag anti-human/NHP IgG antibody (cat no# D20JL-6, lot no# W0019029S, MSD), at 1/1,000 dilution in Blocker™ Casein in PBS was added to each well and plates were then incubated for 1-1.5 hours at room temperature on an orbital shaker. Plates were washed 3X with 1X washing solution and 150 μL of 2X read Buffer (cat# R92TC-3, MSD) was added to each well. Plates were read immediately on an MSD instrument and the STI-2020 standard curve was used to calculate the concentration of antibody present in serum, lung lavage, and organ lysate materials.

### Biodistribution Study

Female CD-1-IGS (strain code #022) were obtained from Charles River at 6-8 weeks of age. For intravenous injection of STI-2020, 100 μL of antibody diluted in 1X HBSS was administered retro-orbitally to anesthetized animals. For intranasal injections, antibody was diluted in 1X HBSS and administered by inhalation into the nose of anesthetized animal in a total volume of 20 μL using a pipette tip. Organs, blood, and lung lavage samples were collected 24 hours post-antibody administration. Blood was collected by retro-orbital bleeding and then transferred to Microvette 200 Z-Gel tubes (Cat no# 20.1291, lot# 8071211, SARSTEDT). Tubes were then centrifuged at 10,000 x g for 5 minutes at room temperature. Serum was transferred into 1.5 mL tubes and stored at −80°C. Lung lavage samples were collected following insertion of a 20G x 1-inch catheter (Angiocath Autoguard, Ref# 381702, lot# 6063946, Becton Dickinson) into the trachea. A volume of 0.8 mL of PBS was drawn into a syringe, placed into the open end of the catheter, and slowly injected and aspirated 4 times. The syringe was removed from the catheter, and the recovered lavage fluid was transferred into 1.5 mL tubes and kept on ice. Lavage samples were centrifuged at 800 × *g* for 10 min at 4°C. Supernatants were collected, transferred to fresh 1.5 mL tubes, and stored at −80°C. Total spleen, total large intestine, and 150 to 400 mg of lungs and small intestine were suspended in 300 μL of PBS in pre-filled 2.0 mL tubes containing zirconium beads (cat no# 155-40945, Spectrum). Tubes were processed in a BeadBug-6 homogenizer at a speed setting of 3000 and a 30 second cycle time for four cycles with a 30-second break after each cycle. Tissue homogenates were centrifuged at 15,000 rpm for 15 minutes at 4°C. Homogenate supernatants were then transferred into 1.5 mL tubes and stored at −80°C. STI-2020 antibody levels in each sample were quantified using the antibody detection ELISA method. Statistical significance was determined using the Welch’s t-test. This study was reviewed and accepted by the animal study review committee (SRC) and conducted in accordance with IACUC guidelines.

### Pharmacokinetic Study

Female CD-1-IGS (strain code #022) were obtained from Charles River Laboratories at 6-8 weeks of age. STI-2020 dissolved in intranasal formulation buffer was administered as described for the IN biodistribution study. Lungs and blood were collected from 3 mice at each of the following timepoints: 10 min, 1.5 h, 6 h, 24 h, 72 h, 96 h, 168 h, 240 h, and 336 h. Serum and lung tissue samples were collected as described for the biodistribution study. STI-2020 antibody levels in each sample were quantified using the antibody detection ELISA method. Pharmacokinetic analysis of the collected ELISA data was performed with the Phoenix WiNnonlin suite of software (version 6.4, Certara) using a non-compartmental approach consistent with an IN bolus route of administration. Statistical significance was determined using the Welch’s t-test. This study was reviewed and accepted by the animal study review committee (SRC) and conducted in accordance with IACUC guidelines.

### Hamster challenge experiments

Female Syrian golden hamsters were obtained from Charles River Laboratories at 6 weeks of age. Hamsters were inoculated IN with 5×10^4^ TCID_50_ of SARS-CoV-2 in 100 μL of sterile PBS on day 0. Antibody treatments were administered IV with monoclonal antibodies (mAbs) against SARS-CoV-2 Spike, or isotype control mAb in up to 350 μL of formulation buffer to anesthetized animals at 12 hours-post inoculation. For intranasal delivery of these antibodies, 100 μL of formulated material was introduced directlΨ into the nares and inhaled by anesthetized animals. Animals were monitored for illness and mortality for 9 days post-inoculation and clinical observations were recorded daily. Body weights and temperatures were recorded at least once daily throughout the experiment. Average % weight change on each experimental day was compared with the isotype control mAb-treated group using 2-way ANOVA followed by Fisher’s LSD test. All animals were housed in animal biosafety level-2 (ABSL-2) and ABSL-3 facilities in Galveston National Laboratory at the University of Texas Medical Branch. All animal studies were reviewed and approved by the Institutional Animal Care and Use Committee at the University of Texas Medical Branch and were conducted according to the National Institutes of Health guidelines.

## RESULTS

### Biodistribution of IV or IN-administered STI-2020 in CD-1 mice

Biodistribution studies of STI-2020 delivered by either the intravenous or intranasal route were carried out in CD-1 mice. Twenty-four hours following administration of a single antibody dose, samples of serum, lung lavage, and tissues including spleen, lung, small intestine, and large intestine were obtained from each of 5 treated mice at each dose level. Samples were processed and mAb levels were quantified using a human antibody detection ELISA readout. Following IV treatment at a dose level of 0.5 mg/kg, STI-2020 was detected in the serum, spleen, lungs, small intestine, and large intestine. Detected levels in the serum at 0.5 mg/kg dose averaged 4.5 μg/mL, while STI-2020 was present at average concentrations less than 0.01 μg/mL in lung lavage material at each of the IV doses tested. Upon processing of lung tissue, antibody was detected at a mean concentration of 0.2 ng/mg of tissue in the 0.5 mg/kg IV dose group. Antibody biodistribution in lung tissue at the 0.05 mg/kg dose level tracked with the 10-fold decrease in the administered dose and STI-2020 was undetectable in the lungs at the lowest dose, 0.005 mg/kg. Antibody levels in the spleen reached a similar average concentration at 24 hours to that seen in lung tissue (0.1 vs 0.2 ng/mg of tissue, respectively). Antibody was detectable in both the small and large intestines at the highest dose level, with similar average concentrations at 24 hours of 0.04 and 0.03 ng/mg of tissue, respectively.

Following administration of STI-2020 by the intranasal route, the concentration of antibody in the serum at 24 h reached an average value of 0.21 μg/mL at the 2.5 mg dose level and was measured at an average value of 0.08 μg/mL in the 0.5 mg/kg dose group. As compared to IV treated animals at the 0.5 mg/kg dose, STI-2020 administered intranasally resulted in a 30-fold lower concentration of antibody in serum. In contrast, STI-2020 concentrations in lung lavage samples following IN dosing reached average concentrations of 2.7 μg/mL in the 2.5 mg/kg dose group, and 1.1 μg/mL in animals dosed at 0.5 mg/kg. In 4 of 5 treated mice, lung lavage STI-2020 levels following IN administration of antibody at the 0.5 mg/kg dose level were elevated between 6 to 37-fold over those observed in lung lavage following administration of an equivalent IV dose. A single mouse in the IN 0.5 mg/kg group displayed much higher lung lavage distribution of antibody, a likely indication of the variability in delivery efficiency in this experiment. STI-2020 was detected in lung tissue samples following an IN dose of 0.5 mg/kg at average concentrations similar to those recorded in IV-treated animals at the same dose level. At a dose of 2.5 mg/kg IN, the average concentration in the lung was measured at 0.91 μg/mg of tissue. Besides the antibody detected in lung tissue, STI-2020 levels in spleen, small and large intestine at all IN dose levels tested did not rise to levels above background in the antibody detection ELISA.

### Pharmacokinetic Studies of STI-2020 administered IN to CD-1 mice

To characterize STI-2020 pharmacokinetics following intranasal dosing at 5 mg/kg, antibody levels in CD-1 mouse lung lysates and serum were quantified at designated timepoints spanning a total of 336 h using a human antibody detection ELISA. The concentration of STI-2020 in lung lysates and serum for each individual mouse are shown in Figures 2A and 2B, respectively. There were no quantifiable concentrations of STI-2020 antibody in the pre-dose samples. Following IN administration of STI-2020, the antibody concentration was quantifiable up to 240 and 336 h in the lungs and serum, respectively. We observed a mouse-to-mouse variability at each time point that could be inherent to the delivery method^15^. When using intranasal instillation, the relative distribution between the upper and lower respiratory tract and the gastrointestinal tract is influenced by delivery volume and level of anesthesia. Average antibody concentration in the lung measured 10 minutes after dosing was nearly 70 percent of the maximum antibody concentration (C_max_) measured during the experiment. The C_max_ value of STI-2020 in the lungs was measured at 1.5 hours post-administration at a value of 43 μg/mL. In the lungs, an apparent terminal half-life (T_1/2_) of 32.21 hours was measured when analyzed between 0.15 and 240 h (Table 1). Under these conditions the R^2^ value equaled 0.932, however when the data were analyzed between 0.15 and 168 hours the R^2^ value increased to 0.987 but the T_1/2_ dropped to 25.07 hours for the lung samples. Kinetics of STI-2020 exposure in the lungs following intranasal administration was accompanied by a slower kinetic of detectable antibody in the serum of treated mice (Figure 2C). Antibody was first detected in the serum at 6 hours post-administration and the C_max_ of 871 ng/mL was detected at the 240-h timepoint (T_max_). Serum antibody concentrations were within 90% of the recorded C_max_ by the 24-h timepoint. Antibody levels remained relatively constant in serum over the period spanning 24-240 h, which is in keeping with the calculated STI-2020 serum half-life observed following IV administration of 240 hours in mice (data not shown). The total systemic exposure (AUC_last_) was significantly higher in the lungs than in the serum of treated mice (AUC_last_ were 1,861,645.8 and 248,675.5 h*ng/mL respectively, Table 1). Although some mice may have generated anti-drug antibody (ADA) during the course of the study the data (measured antibody concentrations and PK profiles) do not suggest that PK parameters were significantly influenced by immunogenicity.

**Figure 1.**
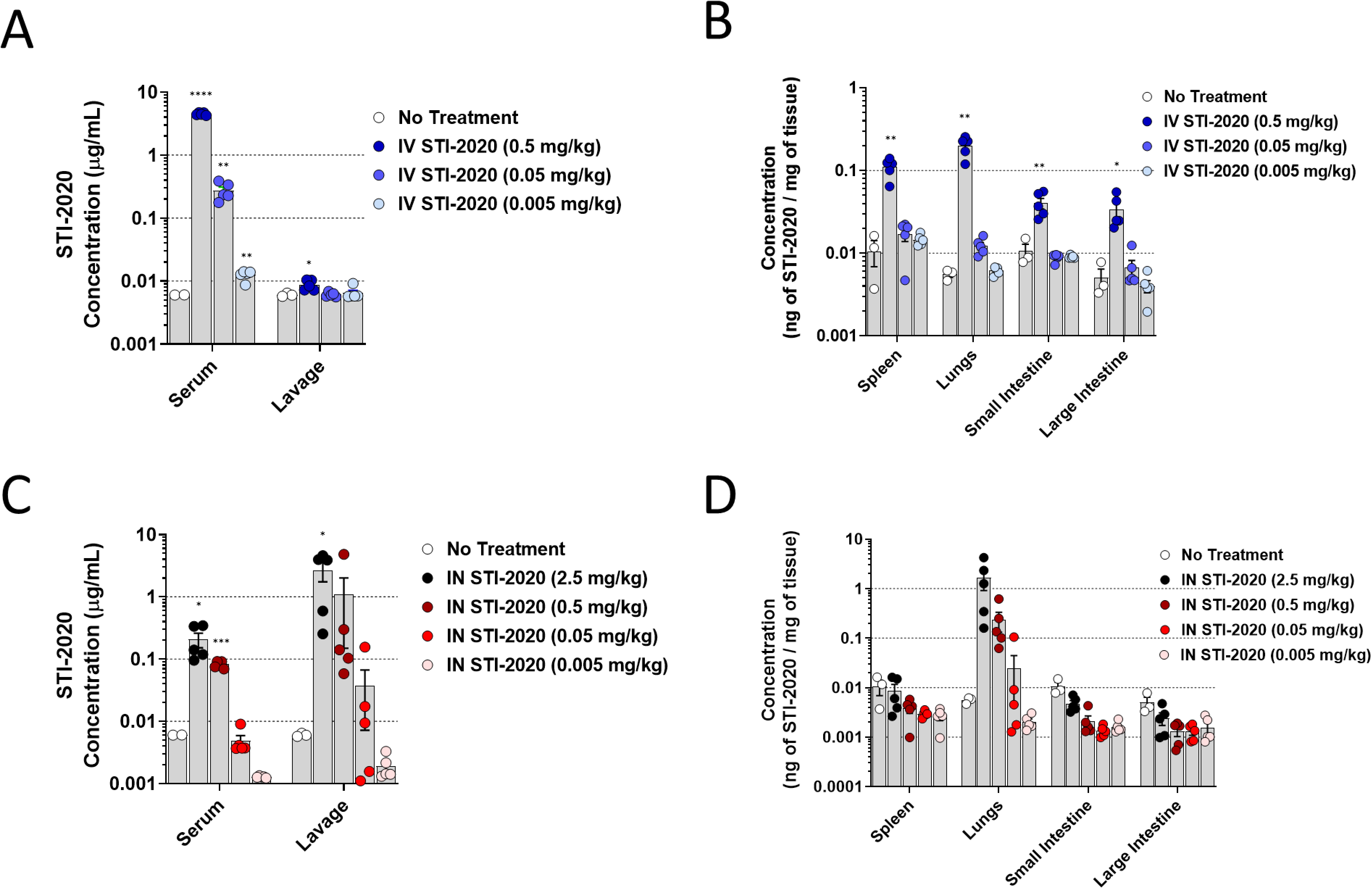
Biodistribution of STI-2020 following IV or IN Administration. **(A)** Concentration of STI-2020 in serum and lung lavage collected from female CD-1 mice administered STI-2020 intravenously (IV) at doses of 0.5 mg/kg 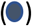, 0.05 mg/kg 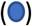, or 0.005 mg/kg 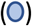 at 24 hours post-administration as compared to samples collected from untreated mice. **(B)** Concentration of STI-2020 in lysates of collected spleens, lungs, small intestines, and large intestines from animals dosed at the same dose levels detailed in panel A. **(C)** Concentration of STI-2020 in serum and lung lavage collected following administration of STI-2020 intranasally (IN) at doses of 2.5 mg/kg 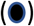, 0.5 mg/kg 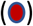, 0.05 mg/kg 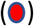, and 0.005 mg/kg 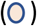 as compared to samples from untreated mice. **(D)** Concentration of STI-2020 in lysates of collected spleens, lungs, small intestines, and large intestines from animals dosed at the same dose levels detailed in panel C. Values represent mean ± SEM (n=3 animals no treatment group, n=5 in treatment groups). Significant differences are denoted by *, P < 0.05; **, P < 0.01; ***, P < 0.001, ****, P < 0.0001.

**Figure 2.**
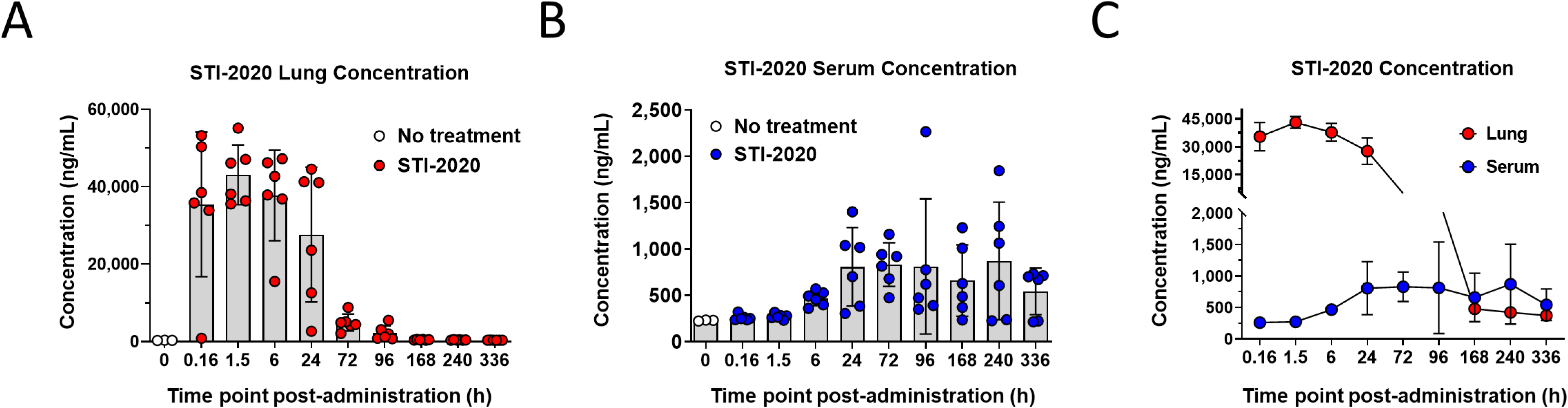
Pharmacokinetics of Intranasally Administered STI-2020 in Mouse Lungs and Serum. **(A)** Concentration of STI-2020 in lung tissue collected from female CD-1 mice administered STI-2020 intranasally (IN) at a dose of 5 mg/kg. Samples from treated mice 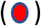 were collected at the indicated timepoint post-administration and STI-2020 antibody concentrations were quantified by ELISA and compared to samples collected from untreated mice. **(B)** Concentration of STI-2020 in serum isolated from female CD-1 mice administered STI-2020 intranasally (IN) at a dose of 5 mg/kg. Serum samples were collected from treated mice 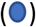 and compared to samples from untreated mice. **(C)** Overlay of STI-2020 concentrations in lung tissue vs. serum following IN administration of a 5mg/kg dose. Values represent mean ± SD (n=3 animals no treatment group, n=6 per time point in treatment groups).

**Figure 3.**
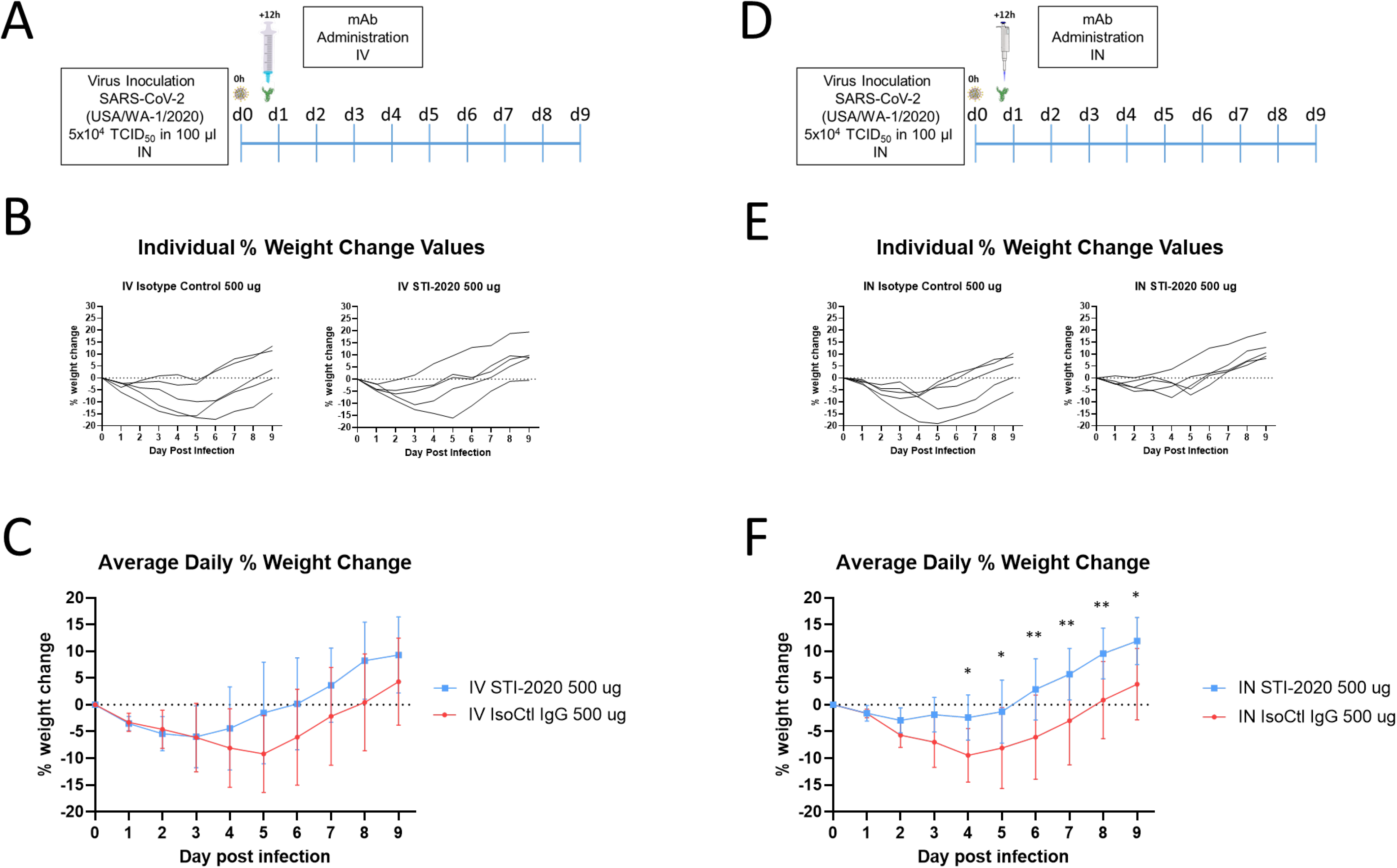
Protective Efficacy of STI-2020 Administered Intravenously or Intranasally in the Syrian Golden Hamster Model of COVID-19. **(A)** Female hamsters were inoculated with SARS-CoV-2 WA-1 isolate on day Twelve hours post-infection, animals (n=5 per group) were administered a single intravenous dose of Control IgG (500 μg, red circles) or STI-2020 (500 μg, blue squares). Daily weight changes from day 0 to day 10 were recorded and **(B)** plotted for each individual animal. **(C)** Average % daily weight change ± standard deviation was plotted for each group. **(D)** Female hamsters were infected as described in panel A. Twelve hours post-infection, a single dose of 500 μg STI-2020 or Control IgG was administered intranasally (IN) and daily weight changes were recorded and **(E)** plotted for each individual animal. **(F)** Average % daily weight change ± standard deviation was plotted for each group. Days on which there was a significant difference in average % weight change between 500 μg STI-2020-treated animals (blue squares) and 500 μg Control IgG-treated animals (red circles) are denoted by *, P < 0.05; **, P < 0.01.

**Table 1.** PK Parameters of STl-2020 in Mice by non-compartmental analysis (NCA)

| Time Boundaries (h) | Organ | T<sub>max</sub> (h) | C<sub>max</sub> (ng/mL) | Terminal half-life (h) | AUC<sub>last</sub> (h*ng/mL) | R<sup>2</sup> |
| --- | --- | --- | --- | --- | --- | --- |
| 0.15-240 | Lung | 1.5 | 43073.16 | 25.07 | 1861645.8 | 0.987 |
|  | Serum | 240.0 | 871.02 | 664.99 | 248675.5 | 0.444 |
| 0.15-168 | Lung | 1.5 | 43073.16 | 32.21 | 1861645.8 | 0.932 |
|  | Serum | 240.0 | 871.02 | 664.99 | 248675.5 | 0.444 |

### Treatment using IN or IV administered STI-2020 in a hamster model of COVID-19

Based on the observed kinetics of STI-2020 exposure in the lungs following IN dosing at 5 mg/kg and considering the protective efficacy of the 5 mg/kg IV dose in the Syrian golden hamster model of COVID-19, we chose 5 mg/kg as our IN and IV dose to be administered 12h post-infection in the hamster SARS-CoV-2 disease model. In this manner, we were able to directly compare the degree of disease severity and duration of disease in animals receiving a therapeutic 5 mg/kg dose of STI-2020 or a control IgG1 antibody (IsoCtl) by either the IV or the IN route. Animals were infected with 5×10^4^ TCID_50_ of SARS-CoV-2 intranasally and subsequently treated with STI-2020 administered intravenously or intranasally at 12 h post-infection. Weight change as a percentage of starting weight was recorded and graphed for each animal.

Animals administered a 5 mg/kg dose of STI-2020 intravenously experienced a progression of disease similar to that of IsoCtl-treated animals for the first three days of infection. Based on the day over day average rate of weight change between the two treatment groups, the STI-2020-treated animals showed a slight decrease in the rate of weight loss between day 2 and day 3. By day 4 of infection, the weight loss rates had further separated along this same trend, and animals in the STI-2020 treatment group had begun to gain weight, on average. On day 5 of infection, the day on which the maximum average percentage weight loss in the IsoCtl group was observed (9.2%), the STI-2020-treated animals had already experienced two consecutive days of weight gain (average 2.2 grams/day). Average weight gain between day 3 and day 8 of the experiment between the STI-2020 and the IsoCtl IV-treated groups was 2.8 grams/day and 1.3 grams/day, respectively. Treatment with STI-2020 12h post-infection decreased duration clinical signs of disease by at least 24 h and led to an overall reduction of disease severity, as manifested by an average rate of weight gain double that of IsoCtl-treated animals between day 3 and day 8 of infection among STI-2020 treated animals. Of note, one of the animals in the STI-2020 treatment group exhibited a more profound course of disease than the other four animals in the same treatment group. Larger experimental groups as well as a dose-response study design will be required to better appreciate factors such as variable disease progression prior to antibody treatment that might contribute to this class of outcomes in the therapeutic treatment setting.

Animals treated IN with STI-2020 had a maximum average weight loss of 2.9% of starting weight as compared to a maximum average weight loss level of 9.5% recorded in IN IsoCtl-treated animals. Weight loss is the major clinical sign of disease recorded in our experiments with this model, and, as such, the mild weight loss observed in the STI-2020-treated animals represents a mitigating effect of the IN-administered antibody on the severity of disease. STI-2020-treated animals maintained their average weight over the first four days of infection, while IsoCtl animals steadily lost weight across this timespan. Beginning on day 4 of infection and extending to day 8 of infection, the weight of STI-2020 treated animals was significantly different than that of animals in the IsoCtl-treated group. By day 8, animals treated IN with STI-2020 had reached an average weight 9.6% above that of their average starting weight, while the average day 8 weight of animals in the IsoCtl group was 0.9% of the average day 0 weight. In general, the difference in average group weights at day 8 reflects both a difference in disease severity in the STI-2020-treated animals between day 0 and day 5 of infection, as well as 20% higher average rate of weight gain among STI-2020-treated animals between days 5 and 8 of infection (3.6 grams/day) vs. that in the IsoCtl-treated group (3 grams/day).

## DISCUSSION

Biodistribution of STI-2020 in the lungs and lung lavage material following IN administration of a nasal drop antibody formulation was elevated well above that observed in mice administered an equal dose of STI-2020 by the IV route. Based on this result, and given that STI-2020 IN lung PK study revealed an antibody half-life of approximately 25 h in lung tissue, an experiment to determine the protective efficacy of IN administered STI-2020 in the hamster model as compared to antibody administered IV was undertaken. In addition, treatment was administered at 12 h post-infection, a regimen designed to mimic a COVID-19 therapeutic scenario involving a newly diagnosed or a more advanced state of disease progression in the upper and lower respiratory tract by allowing establishment of virus infection in the upper airway and along the respiratory tract by the virus inoculum. The immediate effects of administering STI-2020 IN were similar to those observed in the initial stages of disease when a 5 mg/kg dose of STI-2020 was administered IV immediately following infection (1h post). In that instance, the maximum average weight loss for STI-2020 animals was 1.9%. This occurred on day 2 of infection and animals weight returned to levels above the starting weight by day 4 of infection. Animals treated at 5 mg/kg IV 1h post-infection continued to gain weight at a steady rate for the remainder of the experiment. In the case of IN treatments at 12 h post-infection, average weight loss was limited to less than 3% in the STI-2020 treatment group, an indication of decreased disease severity when compared to an average weight loss of over 9% in IsoCtl-treated hamsters. While the effects of infection were diminished, average weight in the STI-2020 treatment group did not reach average day 0 weight levels until day 6 of infection, and weight change dynamics in this group suggest a diminished pathogenic process may have been acting on these animals until day 4 of infection.

Treatment of animals IV at 12 h post infection resulted in a maximum average weight loss of nearly 6%, which occurred on day 3 of infection. As such, IV-treated animals experienced an early course of disease that was indistinguishable from IsoCtl-treated animals and marginally more severe than the disease seen in IN-treated animals across that timespan. Once IV-administered antibody began to demonstrate antiviral efficacy, animals gained weight at a rate similar to that seen after day 5 of infection in IsoCtl-treated animals. By day 5 in the IV STI-2020-treated group, animals had, on average, returned to their day 0 weight and continued to steadily gain weight until the experiment was ended on day 9. The improvements in early disease mitigation following IN dosing of hamsters may reflect the corresponding increases in the measured concentration of STI-2020 in lung lavage samples from mice in our biodistribution studies. Extravascular antibody in the lung may provide more acute protection against disease progression in the lung parenchyma than antibody present in the lung vasculature at early stages of disease. Combining the mitigating effects of IN-administered antibody on early disease severity with the effects of IV antibody dosing on disease duration may prove to be a regimen that maximizes the therapeutic effects of neutralizing antibodies on respiratory symptoms associated with COVID-19. Effects of varying the virus challenge dose and the single or combined IN and IV STI-2020 dose levels on disease duration and severity will be measured in future experiments using the hamster model.

## Author Contributions

R.A., H.J., S.P., M.B., Y.F, D.B., and J.M. conceptualized and designed experiments. J.M., A.S., R.L., A.L., L.R.N., D.L., J.K., I.R., R.S., J.T.M performed experiments and Y.F., J.M., A.S., R.L., L.K., D.B., H.J., S.P., R.A. analyzed data. R.A., D.B., J.M., S.P. wrote the paper.

